# Distractor suppression operates exclusively in retinotopic coordinates

**DOI:** 10.1101/2024.02.01.578407

**Authors:** Yayla A. Ilksoy, Dirk van Moorselaar, Benchi Wang, Sander A. Los, Jan Theeuwes

**Author notes:** **Corresponding author:** Correspondence should be addressed to Yayla Ilksoy, Department of Experimental and Applied Psychology, Vrije Universiteit Amsterdam, Van der Boechorststraat 7, 1081 BT Amsterdam, The Netherlands. **Contact information remaining authors:**. ^1^The authors declare no competing financial interests.

## Abstract

Our attention is influenced by past experiences, and recent studies have shown that individuals learn to extract statistical regularities in the environment, resulting in attentional suppression of locations that are likely to contain a distractor (high-probability location). However, little is known as to whether this learned suppression operates in retinotopic (relative to the eyes) or spatiotopic (relative to the world) coordinates. In the current study, two circular search arrays were presented side by side. Participants learned the high-probability location from a learning array presented on one side of the display (e.g., left). After several trials, participants shifted their gaze to the center of the the test array (e.g., located on the right side) in which all locations were equally likely to contain the distractor. Due to the saccade, the test array contained both a spatiotopic and a retinotopic matching location relative to the original high-probability location. The current findings show that, following saccadic eye movements, the learned suppression remained in retinotopic coordinates only, with no measurable transfer to spatiotopic coordinates. Even in a rich environment, attentional suppression still operated exclusively in retinotopic coordinates. We speculate that learned suppression may be resolved by changing synaptic weights in early visual areas.

**Significance statement:** In our daily lives, attention is shaped by past experiences, guiding us to suppress locations that are likely to contain distractions. While this phenomenon has been studied extensively with static search displays, the real world is dynamic - we are constantly moving our eyes. This study addressed this issue by investigating what happens when we learn to suppress a likely distractor location while making eye movements. Do we suppress the same location in space (spatiotopic), or does the learned suppression persist relative to our eyes (retinotopic)? The current findings provide clear evidence of suppression in retinotopic coordinates only.

## Introduction

Where and what we attend is not only influenced by the dynamics of sensory input (bottom– up) and our current goal states (top–down or behavioral relevance) but also heavily influenced by what we have encountered in the past. One example of selection biases implemented by selection history comes from recent studies demonstrating that human observers can learn to extract statistical regularities in the environment resulting in attentional suppression of locations that are likely to contain a distractor, effectively reducing the amount of distraction (Wang & Theeuwes, 2018a, 2018b, 2018c). The general idea is that just like top-down, and bottom-up attention, selection history feeds into an integrated priority (salience) map, ultimately resulting in a winner-take-all competition that determines the allocation of covert and overt attention (Theeuwes, 2019; Theeuwes et al., 2022). The notion of learning-induced plasticity within the spatial priority map is important, as it can explain how lingering biases from former attentional deployments come about. While it is generally agreed that spatial priority maps are topographically organized maps of the external visual world (e.g., Bisley & Goldberg, 2010; Fecteau & Munoz, 2006; Thompson & Bichot, 2005), it remains largely unclear how the “external world” is represented within these maps. As such it remains unclear whether suppression effects due to statistical learning, which is thought to operate via changes of weights within the spatial priority map, operate in retinotopic (relative to the eyes) or spatiotopic (relative to the world) coordinates.

Researchers have identified potential spatial priority map candidates among various brain regions, such as the superior colliculus (Bisley, 2011; Krauzlis et al., 2013; Noudoost et al., 2010; Wurtz et al., 2011a), caudate nucleus (Kim & Hikosaka, 2013; Yamamoto et al., 2012), and regions in the posterior parietal (Bisley & Goldberg, 2010; e.g., lateral intraparietal cortex), frontal (Thompson et al., 2005; Thompson & Bichot, 2005; e.g., frontal eye field) and visual cortices (Li, 2002; Mazer & Gallant, 2003; Zhang et al., 2012). Regardless of whether these regions are cortical or subcortical, it is generally accepted that retinotopy is preserved throughout the brain, suggesting that priority maps are retinotopically organized. Nevertheless, a topographical representation would be more appropriate as it reflects the external visual world upon which we act (e.g., Bisley & Goldberg, 2010; Fecteau & Munoz, 2006; Thompson & Bichot, 2005). If a location is relevant for selection or requires suppression, it makes sense to connect it to external world coordinates rather than retinal location. In line with both views, previous studies have shown that both endogenous attention (Golomb et al., 2008, 2010) and exogenous attention (Mathôt & Theeuwes, 2010a, 2010b) rely on retinotopic maps, which are progressively transformed into spatiotopic maps following saccades. Moreover, recent studies by van Moorselaar & Theeuwes (2023, 2024) showed that people can learn to prioritize a likely target location within objects, irrespective of the object’s orientation in space. This implies that statistical learning is not necessarily limited to retinotopic maps. However, it is still unclear whether history-driven suppression effects persist in retinotopic coordinates or transfer to spatiotopic coordinates after eye movements.

In the present study, we adopted the additional singleton task used by Wang and Theeuwes (2018a) in which the distractor singleton was presented more often in one location than in all other locations. Critically, this regularity was only present when participants were performing the task at one side of the display (labelled as “learning array”). After performing several trials within this learning array (e.g., on the left side), participants shifted their gaze to another display (e.g., the one on the right) and continued the search task, but now without any statistical regularities included (labelled as “test array”). Due to the saccadic eye movement towards the test location, it contained both a spatiotopic matching and a retinotopic matching location relative to the suppressed location in the learning array. The question then was whether the learned suppression within the learning array would stay in retinotopic coordinates, transfer to spatiotopic coordinates, or relies on both coordinate systems.

## Experiment 1

### Methods

#### Participants

To determine an adequate sample size, we conducted an a priori power analysis using simulated data. Previous studies with similar designs (Wang & Theeuwes, 2018a, 2018b) identified a difference of approximately 50 ms between high probability and low probability locations, with an average reaction time of around 800 ms. In our study, we anticipate that the critical effect between the retinotopic, spatiotopic, and low probability locations will be attenuated compared to previous studies, as the learning effect needs to transfer from the learning array to the test array.

Based on these considerations, we expected a slope of 20 ms (β) across the retinotopic, LP, and spatiotopic conditions. We simulated data for 24 participants, with 35 trials per condition, and included the same fixed-effects and random-effects structure as mentioned in the statistical analysis. In addition to a slope for the Distractor condition, we also included slopes for target location priming and distractor location priming. Previous findings (Maljkovic & Nakayama, 1996) suggest that participants are approximately 10 ms faster when the target appears in the same location as the previous trial. We expected a similar but smaller trend for distractor location priming (5 ms). For the other fixed effects, we did not have any theoretical expectations, so we included those fixed effects with a slope of zero.

The simulation-based power analysis, performed using the *simr* package of Green & Macleod (2015) in R (R Core Team, 2018), indicated that with 24 participants, our study would have sufficient power to detect the specified effect size for the Distractor condition with 78.2% power (95% confidence interval, CI [75.51%, 80.72%] in 1,000 simulations) and an alpha level of 0.05.

Twenty-four adults (20 women, mean age: 23.8 years old) were recruited for money compensation or course credits. They all signed informed consent before the study and reported normal or corrected-to-normal visual acuity. The Ethical Review Committee of the Faculty of Behavioral and Movement Sciences of the Vrije Universiteit Amsterdam approved the present study.

#### Apparatus and stimuli

Participants were tested in a dimly lit laboratory, with their chin held on a chinrest located 70 cm away from a 24-in. liquid crystal display (LCD) color monitor. The experiment was created in *OpenSesame* (Mathôt et al., 2012) and run on a Dell Precision 3640 computer. An eye-tracker (EyeLink 1,000) was used to monitor participants’ eye movements and the sampling rate was set to 1,000 Hz.

A modified additional singleton paradigm was adopted. The visual search display consisted of six discrete stimuli with different shapes (one circle vs. five diamonds, or vice versa), each containing a vertical or horizontal gray line (0.2° × 1°) inside (see Figure 1). The stimuli were presented on an imaginary circle with a radius of 3.5°, centered at the fixation (a white cross measuring 0.5° × 0.5°) against a black background (RGB: 0/0/0). The radius of the circle stimuli was 1°, the diamond stimuli were subtended by 1.55° × 1.55°, and each had a red (RGB: 253/34/34) or green (RGB: 90/174/20) outline.

**Figure 1.**
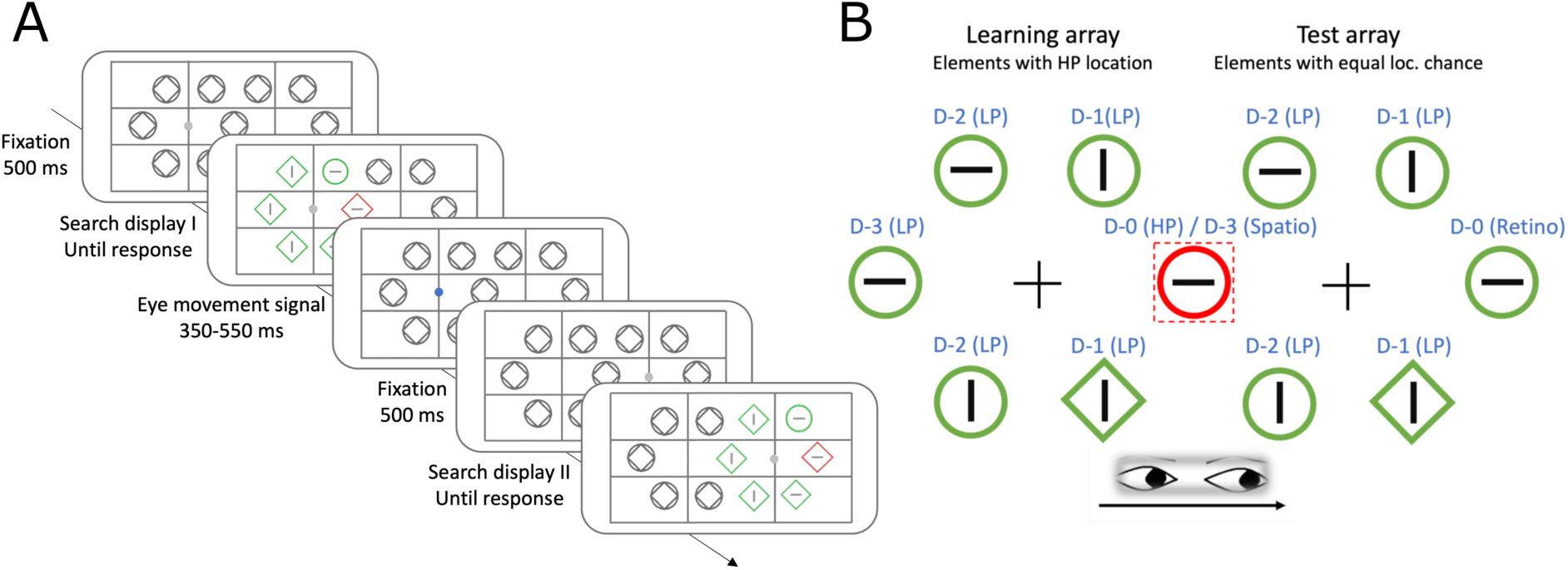
Stimuli and design. *Note.* **(A)** An example of a trial sequence. In this example, the fixation switches from left (Search display I) to right (Search display II). The gridlines and placeholders were introduced in Experiment 2 (see Methods section of Experiment 2a) and were not present in Experiment 1. The placeholders were presented to participants at all possible stimulus locations. **(B)** Possible stimulus locations. The high-probability distractor location was always in the center of the screen (see red striped square marking). The “D” labels represent the distance to the (mapped) HP location. In the test array, location D-3 (Spatio) represents the spatiotopic location and location D-0 (Retino) the retinotopic location. The location of the learning array (left or right), and consequently the position of the HP location within the learning array, was counterbalanced across participants.

#### Experimental design

Every trial started with a fixation cross that remained visible throughout the trial. The fixation cross was presented on the horizontal centerline at 3.5° either left or right of the center of the screen. After 500 ms, a search array was presented and centered at the fixation cross for 2000 ms or until response. Participants searched for one circle (target) among five diamonds (distractors) or vice versa and responded to the orientation of the line segment as fast as possible, by pressing the ‘up’ arrow key for vertical and the ‘left’ arrow key for horizontal with their right hand. The inter-trial interval (ITI) was randomly chosen from 350 to 550 ms.

A target was presented in each trial with an equal probability of being a circle or diamond. A uniquely colored distractor singleton was present in 66.7% of the trials, with the same shape as the other distractors but with a different color (red or green with an equal probability). All conditions were randomized within each block. For each search array, the target could appear at each of the six locations. Importantly, two types of search arrays were presented: a learning and test array. For the learning array, in the distractor present condition, the distractor singleton had a high proportion of 63% to be presented at the center of the display (e.g., the furthest right location of the left search array or the furthest left location of the right search array). This location is called the high-probability (HP) location. Each of the other locations independently had a low proportion of 7.4% to contain a distractor singleton (low-probability location). For the test array, all the locations contained a distractor singleton equally often (16.7% in distractor-present trials). The target location was determined randomly on each trial.

The experiment consisted of six blocks of 250 trials each. The first two blocks only presented the learning array on one side of the display. The position of the learning array (left or right), and consequently the position of the HP location within the learning array, was counterbalanced across participants. After the first two blocks, the learning array alternated with the test array, which was presented on the opposite side of the display. After a randomly selected sequence of 8, 9, or 10 consecutive trials for the learning array or 4 or 5 consecutive trials for the test array, a white dot appeared at the previous fixation location during the ITI period. Following this, participants had to immediately move their eyes to the other fixation on the opposite side of the display to perform the search task for the other search array. Crucially, the location at the center of the screen was shared by the learning and test array: This was the HP location of the learning array and the spatiotopic location of the test array. The retinotopic location was at the opposite side of the test array (see Figure 1B for an illustration). In blocks three to six, the learning and test arrays were presented in 165 and 85 trials, respectively.

There were two practice sessions before the first block of the experiment: one practice session of 15 trials with only the learning array that remained in the same location (as in the first two blocks of the experiment) and one practice session of 40 trials that alternated between the learning and test array (as in block three to six of the experiment). If participants did not achieve more than 70% accuracy or were not faster than 1100 ms on average in the practice sessions, they had to repeat the session. If participants did not respond or made an erroneous response, a warning message was presented. At the end of the experiment participants were asked whether they noticed the statistical regularities (subjective measure) and on which location within the array they thought the HP location was (objective measure). Notably, these questions were interspersed with unrelated questions that were included to avoid influencing responses to the study-related questions.

Participants were instructed to fixate on the fixation cross in every trial. A warning sound was played if eyes deviated from fixation (see Data analysis for further details). Before every block, the eye tracker was calibrated, and an automatic drift check was performed at the beginning of every 10 trials.

#### Statistical analysis

Participants with an average accuracy below 2.5 standard deviation from the overall accuracy were excluded as outliers and replaced. Trials on which the response times (RTs) were shorter than 200 ms and trials on which RTs were shorter or longer than 2.5 standard deviations from the average RT per array per block per participant were excluded from analyses. Subsequently, participants with an average RT longer than 2.5 standard deviations of the group mean were excluded as outliers and replaced. Trials in which eyes deviated from fixation were also excluded. Eye deviations were determined by identifying instances where fixations extended beyond 2.5° from the fixation cross for more than 75 ms (Golomb et al., 2008; Mathôt & Theeuwes, 2010a; Talsma et al., 2013). For RT analyses, only trials with a correct response were included.

Previous work by Wang and Theeuwes (Wang & Theeuwes, 2018a, 2018b) showed that not only the HP location but also its nearby locations were suppressed by learning statistical regularities. In other words, suppression was characterized by a spatial gradient centered at the HP location such that differences in distractor processing were most pronounced between the HP and the opposite location in the display. Since the retinotopic and spatiotopic location were on opposite sides of the display in the current set-up (see figure 2), we expected that retinotopic suppression would result in a gradient from the retinotopic location towards the spatiotopic location (see figure 2A). Conversely, if the suppression effect is spatiotopic, we expected the gradient to occur in the opposite direction (see figure 2b).

**Figure 2.**
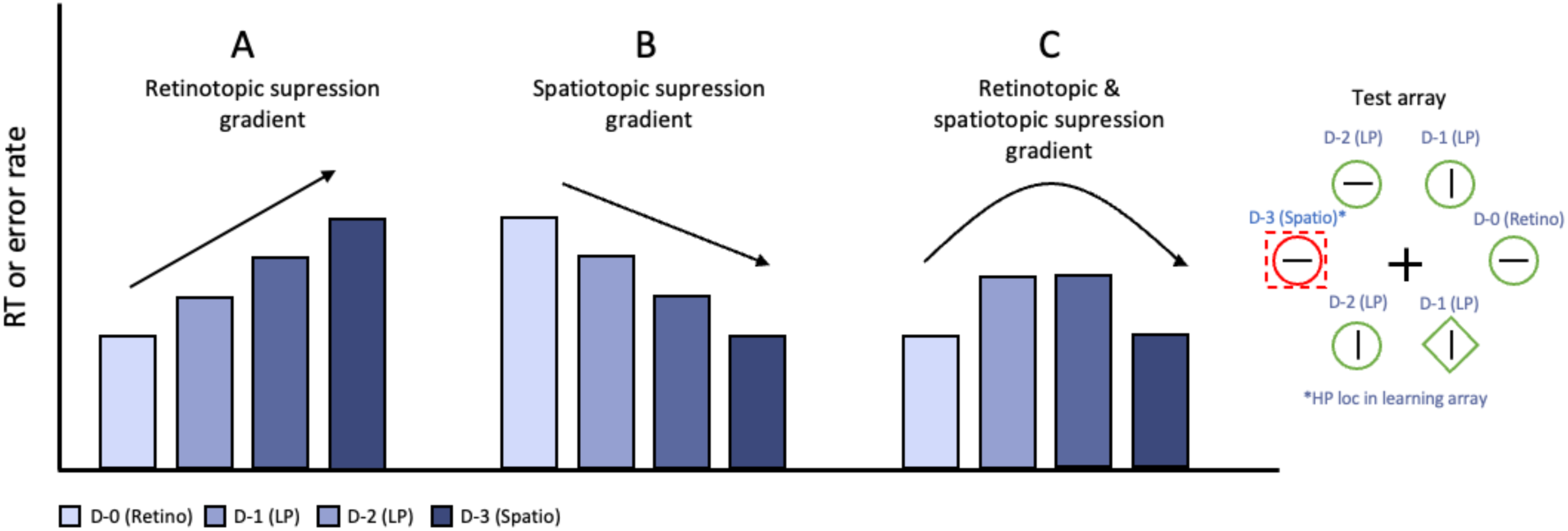
The three hypothesized outcomes in the test array. *Note.* Each bar represents the mean RT or error rate when the distractor is presented at a certain distractor location (D-0, D-1, D-2, D-3). **(A)** An increasing slope across retinotopic, LP and spatiotopic locations suggests a retinotopic centered suppression gradient. **(B)** A decreasing slope across retinotopic, LP and spatiotopic locations suggests a spatiotopic centered suppression gradient. **(C)** A negative parabola across retinotopic, LP and spatiotopic locations suggests both a retinotopic and a spatiotopic centered suppression gradient.

The main analysis was separated into two analytical approaches. First, to ascertain that observers learned to suppress the HP location, learning array RTs and error rates were analyzed using repeated-measures analysis of variance (ANOVAs) followed by planned comparisons with paired-sample t-tests. Where sphericity was violated, Greenhouse-Geiser corrected p-values are reported. For multiple comparisons, p-values were adjusted using the Holm-Bonferroni method. To then determine whether the learned attentional bias, once established, transferred to retinotopic or spatiotopic coordinates, the analysis of the test array included only data from those participants who exhibited visual statistical learning effect in the learning array (see Supplementary Table 1 for parallel analyses involving all participants, which yielded the same overall pattern of results). In contrast to the conventional ANOVA approach here we relied on linear mixed model (LMM) and generalized mixed model (GLMM) approaches for the RT and error rate analyses respectively, where the data is not averaged but instead grouped per participant. For the present purposes, this approach has two main advantages. First, a range of continuous and categorical variables can be added to a single model such that rather than excluding large subsets of data in a series of control analyses, which inevitably reduces power (Brysbaert & Stevens, 2018), various control factors that could potentially modulate the effect of interest can be simultaneously included allowing for a more refined control. Specifically, in all adopted models Distractor condition was incorporated into the fixed-effects structure as an ordered factor. This factor consisted of four levels that described the distance to the mapped HP location in the learning array (cf. Figure 1): D-0 (Retino), D-1 (LP), D-2 (LP), and D-3 (Spatio).

In addition to the main effect of interest, the following factors were entered into the fixed-effects structure: intertrial location distractor and target priming (i.e., whether the position of a distractor or target repeated from one trial to the next; yes, no), array switch (i.e., whether the array position was the same as on the previous trial or had switched; yes, no), target and distractor position (0-5), learning array position (left, right), awareness of the HP location (response to objective measure, see Procedure and design for further details; correct, incorrect), target color (red, green), target shape (circle, diamond) and target line orientation (horizontal, vertical). Second, and most importantly, this approach allowed us to evaluate whether suppression was best characterized by a model resulting from a gradient centered at either the retinotopic or the spatiotopic location, indicative of retinotopic or spatiotopic suppression respectively (see Figure 2A and B), or alternatively by a model in which both retinotopic and spatiotopic suppression exerted their effects simultaneously (see Figure 2C).

For this purpose, the model included a linear, as well as a quadratic coefficient of Distractor condition. The degrees of freedom of all coefficients were estimated using Satterwaite’s method for approximating degrees of freedom and the F statistics, Z-scores and the corresponding p-values were obtained from the *lmerTest* package (Kuznetsova et al., 2017) in R (R Core Team, 2018). All fixed effects were dummy coded. Following guidelines by Barr et al. (2013), a maximal random effects structure justified by design was included. The full model included by-participant random intercepts and by-participant random slopes for distractor condition, distractor and target position, intertrial location distractor and target priming, array switch, target color, target shape and target line orientation. If the model did not converge, we started by removing the random slope for the last factor mentioned above (target line orientation), then continued sequentially (target shape, target color, etc.) until the model converged. The final LMM included by-participants random intercepts and by-participant random slopes for Distractor condition in the random-effects structure whereas the final GLMM only included by-participant intercepts for Distractor condition.

#### Transparancy and openness

All data and analysis codes are available at https://osf.io/ev7zx/. The study’s design and its analysis were not pre-registered. All data was collected in 2023. Despite the participant pool being mostly female undergraduate psychology students, the study’s focus on fundamental cognitive processes suggests limited constraints on generality, as these mechanisms are foundational and likely to generalize across diverse populations. We anticipate consistent results with different stimuli, provided they elicit a pop-out effect from both the distractor and target elements.

### Results

A total of four participants were excluded and replaced: three due to low accuracies (M = 71%), and one because too many trials were excluded due to excessive eye movements (799 trials out of a total of 1500 trials; see Statistical analysis section within the Methods for the exclusion criteria). Exclusion of incorrect responses (7.7%), data trimming (3.3%) and trials with eye movements (10.7%) resulted in an overall loss of 21.7% of the trials for the RT analyses and 14% of the trials for the error rate analyses.

#### Learning array

Before investigating how distractor suppression remaps following a saccade, we first examined to what extent distractor learning took place in the learning array. Repeated-measures ANOVAs with Distractor condition (no distractor, HP location and LP location) as a within-subject factor revealed a reliable main effect on both mean RTs (F (2, 46) = 104.71, *p* < .001, 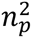 = 0.82; see Figure 3A and 3B) and mean error rates (F (1.55, 35.63) = 32.057, *p* < .001, 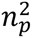 = 0.58; see Figure 3C and 3D). Subsequent planned comparisons showed that relative to no distractor trials, RTs were longer and error rates were higher when the distractor appeared at the HP or LP location (all *t*’s > 5.6 and *p*’s < .001). Critically, RTs were shorter (*t* (23) = 5.27, *p* < .001, *d* = 1.08) and error rates were lower (*t* (23) = 3.90, *p* < .001, *d* = 0.80) when the distractor appeared at the HP location compared to the LP location, indicative of learned attentional suppression at the high probability distractor location. Additionally, it was confirmed that the suppression was characterized by a spatial gradient for both the RTs (linear β = 18.69, SE = 4.77, *t* (14588.11) = 3.92, *p* < .001; quadratic β = -17.53, SE = 7.73, *t* (14588.10) = -2.27, *p* = .023) and the error rates (linear β = 0.17, SE = 0.074, *z* = 2.34, *p* < .05; see Supplementary Materials for figures).

**Figure 3.**
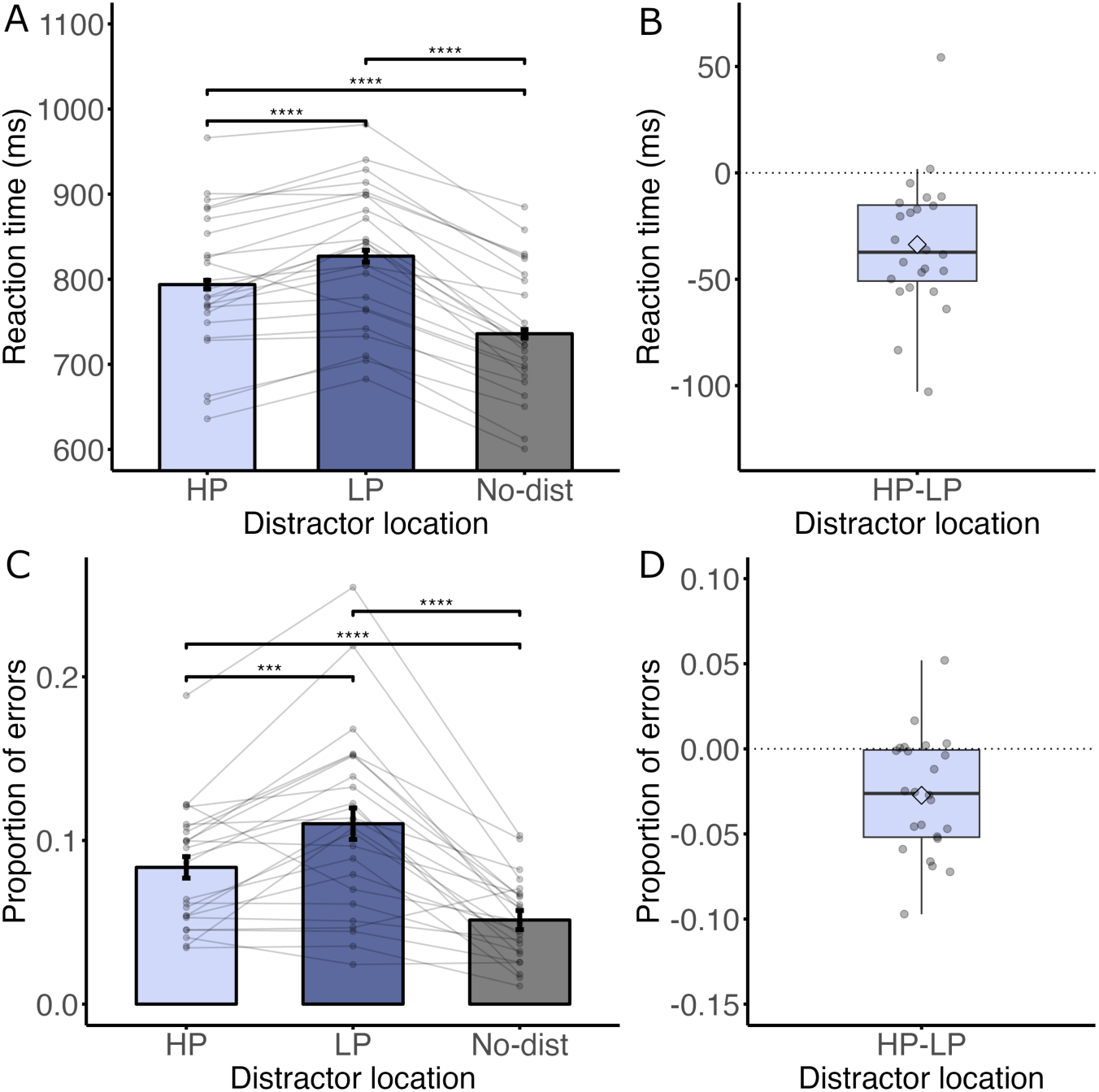
RTs (A and B) and error rates (C and D) in Experiment 1 as a function of distractor location for learning arrays. *Note.* The bars represent the condition means, and each gray dot represents the mean of an individual participant. Error bars represent 95% within-subjects confidence intervals (Morey, 2008). The significance bars represent the planned comparisons with paired-sample t-tests. Significant effects are marked as follows: \**p* < .05, \*\**p* < .01, \*\*\**p* < .001, \*\*\*\**p* < .0001. The diamonds inside the boxplots represent the mean difference scores and the horizontal lines represent the median difference scores. **(A)** RTs in the learning array. The bars show a clear attentional capture effect with longer RTs when the distractor is present. **(B)** The boxplot displays the RT differences between the HP and LP condition in the learning array. Most subjects had shorter RTs when the distractor is presented at the HP location compared to the LP location. **(C)** Error rates in the learning array. The bars show a clear attentional capture effect with higher error rates when the distractor is present. **(D)** The boxplot displays the error rate differences between the HP and LP condition in the learning array. Most subjects have lower error rates when the distractor is presented at the HP location compared to the LP location.

#### Test array

After having established reliable suppression within the learning array, we next set out to establish the dynamics of this learned suppression following a saccade by limiting the analysis to only those participants that showcased learning within the learning array (*N* = 22). We considered three possible scenarios: suppression is retinotopically organized, spatiotopically organized or a combination of both (see Figure 2). As visualized in Figure 4A, progression from the retinotopic towards the spatiotopic location was characterized by a systematic increase in RTs (linear β = 24.30, SE = 8.22, *t* (21.82) = 2.96, *p* < .01), in line with the scenario in Figure 2A (see Supplementary Materials for parallel analyses involving all participants). The error rates followed the same pattern as the RTs, with the gradient approaching significance (linear β = 0.27, SE = 0.15, *z* = 1.81, *p* = .071; quadratic β = -0.35, SE = 0.20, *z* = 1.74, *p* = 0.082; see Figure 4C). Together these findings demonstrate that the observed statistical learning effect did not transfer to spatiotopic coordinates, but instead remained in retinotopic coordinates following a saccade.

**Figure 4.**
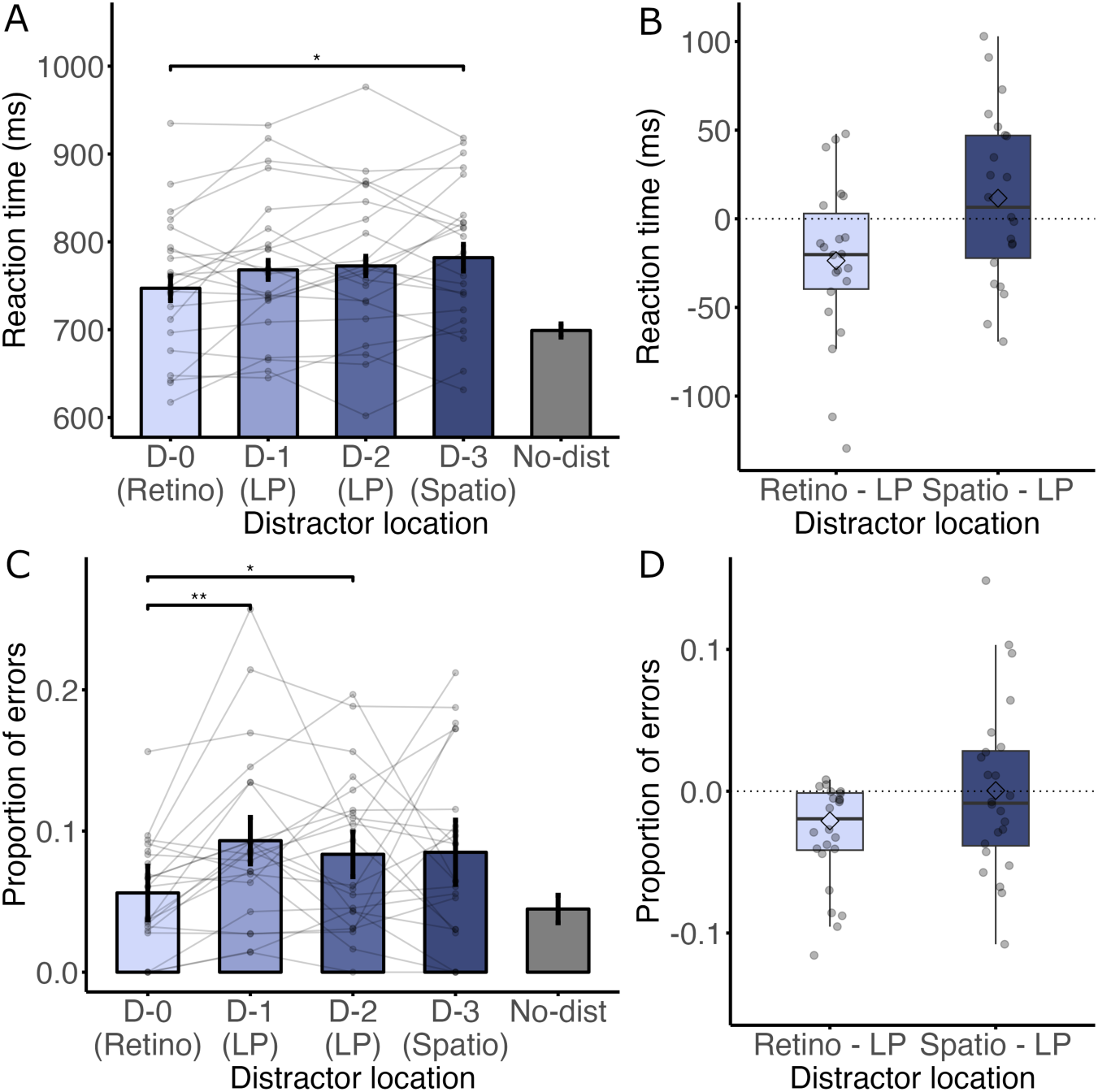
RTs (A and B) and error rates (C and D) in Experiment 1 as a function of distractor location for test arrays. *Note.* While the main analyses used a linear mixed model approach, significance bars based on paired-sample t-tests have been included in the figures to enhance comparability with other studies (see Supplementary Materials for a comprehensive summary of all additional statistical tests). Significance bars between the no distractor condition and each distractor location are omitted in the figure (RT: all *t*’s > 5.3 and *p*’s < .001; Error rate: all *t*’s > 2.9 and *p*’s < .05, except for D-0, *t* = 1.27, *p =* .66). **(A)** RTs in the test array. The bars show a systematic increase in RTs across the retinotopic, LP and spatiotopic locations (linear β = 24.30, SE = 8.22, *t* (21.82) = 2.96, *p* < .01). (B) The boxplot displays the RT differences between the retinotopic and LP locations and the spatiotopic and LP locations. (C) Error rates in the test array. The GLMM fitted a near-significant slope across the distractor locations (linear β = 0.27, SE = 0.15, *z* = 1.81, *p* = .07; quadratic β = -0.35, SE = 0.20, *z* = 1.74, *p* = 0.082) (D) The boxplot displays the error rate differences between the retinotopic and LP locations and the spatiotopic and LP locations.

### Discussion

The current findings show that following a saccadic eye movement, suppression due to statistical learning remained in retinotopic coordinates only, with no measurable transfer to spatiotopic coordinates. While this is an important finding, it should be noted that in the current set-up there were no visual environmental landmarks as the search display was presented on the background of a blank empty screen. Also, with each saccade, the entire display shifted from side to side, making the entire visual field move along with the eye movements. It is therefore possible that the absence of a spatiotopic effect has to do with the absence of any visual landmarks. To that end, a second experiment was conducted with a grid and placeholders in the display to create more structure by introducing visual landmarks (see Figure 1).

## Experiment 2a

### Methods

Experiment 2a was identical to Experiment 1 except for the following changes. The experiment was conducted in an online environment on a JATOS server (Lange et al., 2015). Because the experiment was conducted online, our control over the experimental settings was restricted, and as a result we report the stimuli in terms of pixels instead of visual degrees. The search arrays (search radius was 150 pixels; diamond stimuli were subtended by 56 × 56 pixels, circle stimuli had radius of 45 pixels) were presented inside a gray-colored grid with 4 × 4 horizontal and vertical lines (see Figure 1A). To ensure that the grid remained noticeable, we modified the line thickness three times within each block. At the onset of each block, gridlines were consistently presented with a thickness of 3 pixels. Every 50 trials, the gridline thickness randomly alternated, transitioning between 1, 5, and 7 pixels. This variation was implemented to ensure that the grid did not become visually monotonous. Dark gray placeholders in the form of a circle imposed upon a diamond were presented at all possible stimulus locations, covering both the the learning array and test array positions (11 locations in total). To ensure that the participants maintained fixation effectively before initiating saccades, the stimulus display was presented for only 150 ms, which is a duration that is too short to make any directed eye movements within the search array (Fischer & Ramsperger, 1984; Fischer & Weber, 1993; Heeman et al., 2019). The experiment consisted of five blocks of 200 trials each, with the first block only consisting of arrays presented on one side of the display (either left or right, counterbalanced across participants).

#### Participants

Due to the increased noise in online studies and the challenging nature of Experiment 2a, which could further amplify this noise, we decided to expand the sample size to 50 participants in total. Fifty adults (23 women, mean age: 27.9 years old) were recruited for monetary compensation via the online platform Prolific (www.prolific.co; £10.33).

### Results

Prior to outlier detection (See Statistical analysis section of the Methods of Experiment 1 for the exclusion criteria), seven participants with an average accuracy below 60% (M = 54%), indicative of chance-level performance, were removed and replaced because their low performance reduced the overall mean considerably, which influenced the outlier detection process. Subsequently, a total of five participants were identified as outliers and replaced: one due to low mean accuracy (M = 60%) and four due to long mean RTs (M = 949 ms). Three additional participants were substituted due to stimuli being displayed for over 180 ms (instead of the intended 150 ms) in more than 50% of the trials, attributable to the refresh rate of their personal computers. Exclusion of incorrect responses (18.3%) and data trimming (2.1%) resulted in an overall loss of 20.5% of the trials for the RT analyses and 2.1% of the trials for the error rate analyses. The final models to analyze the RTs and error rates in the test array included by-participant random intercepts and by-participant random slopes in the random-effects structure.

#### Learning array

For the learning array, repeated-measures ANOVAs with Distractor condition (no distractor, HP location and LP location) as within-subjects factor showed a main effect for both mean RTs (*F* (2, 98) = 50.093, *p* < .001, 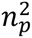 = 0.51) and mean error rates (*F* (2, 98) = 97.32, *p* < .001, 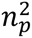 = 0.67). As before, subsequent planned comparisons revealed longer RTs and higher error rates when the distractor was presented at the HP location or LP location compared to the no distractor condition (all *t*’s > 7.1, all *p*’s < .001; see Figure 5A and 5C). Crucially, in comparison to the LP location, RTs were shorter (*t* (49) = 3.26, *p* < .01, *d* = 0.46; see Figure 5B), and error rates were lower (*t* (49) = 4.92, *p* < .001, *d* = 0.70; see Figure 5D) at the HP location, indicating attentional suppression at the high-probability distractor location. As in Experiment 1, it was confirmed that the suppression was characterized by a spatial gradient (RTs: linear β = 12.69, SE = 2.88, t (17635.42) = 4.41, p < .001; Error rates: linear β = 0.21, SE = 0.042, *z* = 5.061, *p* < .001; see Supplementary Materials for figures).

**Figure 5.**
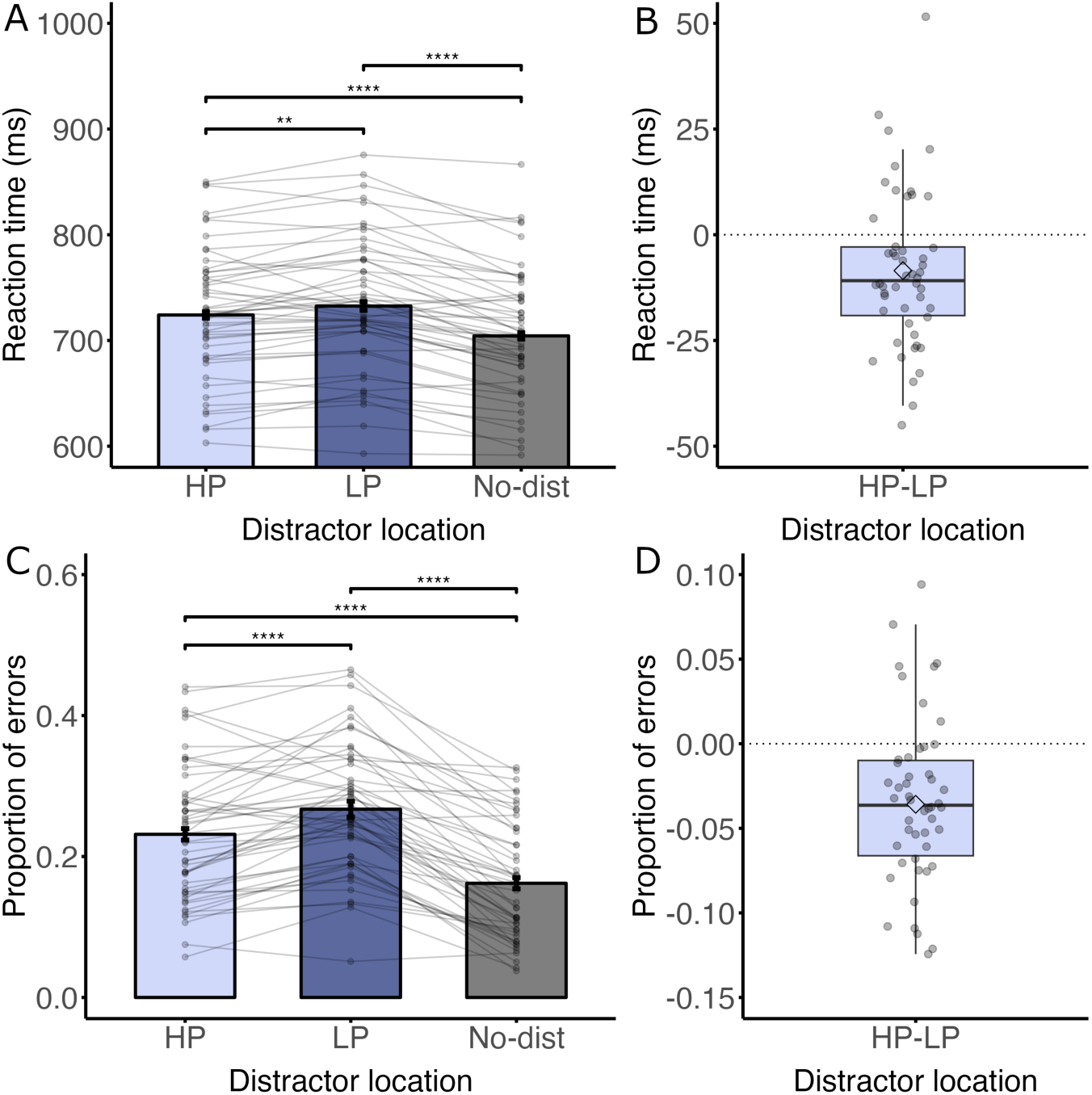
RTs (A and B) and error rates (C and D) in Experiment 2a as a function of distractor location for learning arrays. *Note.* **(A)** RTs in the learning array.The bars show a clear attentional capture effect with longer RTs when the distractor is present. **(B)** The boxplot displays the RT differences between the HP and LP condition in the learning array. Most subjects had shorter RTs when the distractor is presented at the HP location compared to the LP location. **(C)** Error rates in the learning array. The bars show a clear attentional capture effect with higher error rates when the distractor is present. **(D)** The boxplot displays the error rate differences between the HP and LP condition in the learning array. Most subjects have lower error rates when the distractor is presented at the HP location compared to the LP location.

#### Test array

Having established a learned attentional bias in the learning array, we next set out to examine whether that bias continued to persist in retinotopic coordinates after a saccade is made in the presence of environmental landmarks by again including only those participants that demonstrated the hypothesized effect in the learning array (*N* = 38). As visualized in Figure 6A, and unlike the findings of Experiment 1, there was no linear increase from the retinotopic, to the LP to the spatiotopic location (linear β = 6.82, SE = 6.17, *t* (38.44) = 1.10, *p* = .28). Instead, RTs were fastest at the LP location relative to the retinotopic and the spatiotopic location (quadratic β = 26.94, SE = 7.79, *t* (110.30) = 3.46, *p* < .001), a pattern that is inconsistent with any of the models outlined in Figure 2 (see Supplementary Materials for parallel analyses involving all participants). By contrast, error rates did showcase a systematic rise from the retinotopic location towards the spatiotopic location (linear β = 0.21, SE = 0.096, *z* = 2.17, *p* = .030; see Figure 6C). Together, these findings again demonstrate that there was no evidence that learned spatial suppression would be remapped in spatiotopic coordinates following a saccade, not even when visual landmarks provided more visual structure.

**Figure 6.**
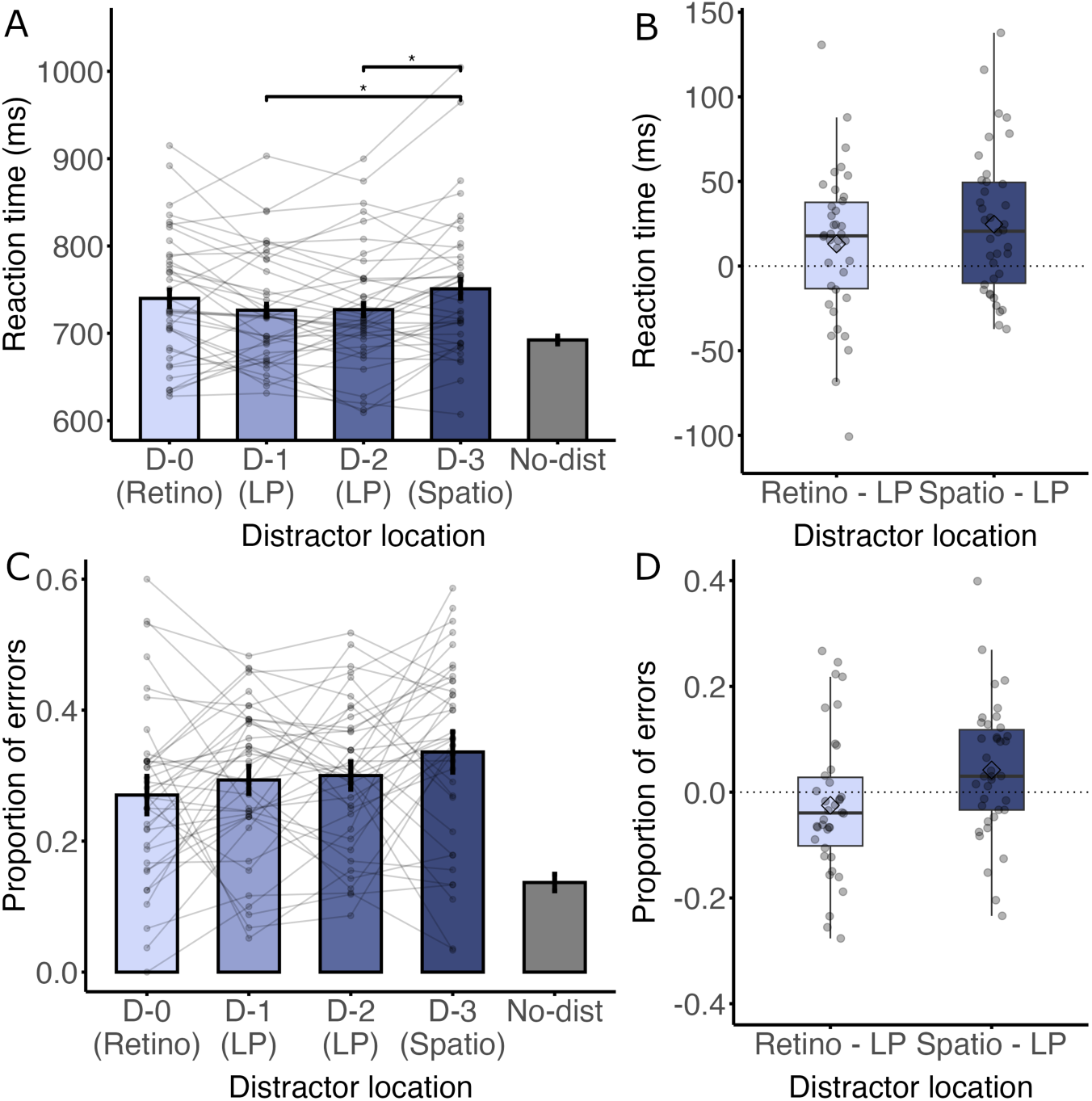
RTs (A and B) and error rates (C and D) in Experiment 2a as a function of distractor location for test arrays. *Note.* Significance bars between the no distractor condition and each distractor location are omitted in the figure (all *t*’s > 5.9 and *p’s* < .001). **(A)** RTs in the test array. The bars show that the RTs are fastest when the distractor is presented at the LP locations (quadratic β = = 26.94, SE = 7.79, *t* (110.3) = 3.46, *p* < .001), which is inconsistent with any of the expected scerarios. **(B)** The boxplot displays the RT differences between the retinotopic and LP location and the spatiotopic and LP location. **(C)** Error rates in the test array. The bars show a systematic increase in error rates across the retinotopic, LP and spatiotopic locations (linear β = 0.21, SE = 0.096, *z* = 2.17, *p* = .030) **(D)** The boxplot displays the error rate differences between the retinotopic and LP location and the spatiotopic and LP location.

### Discussion

In Experiment 2a, we added a grid and placeholders to the search display to impose a spatial reference frame and promote spatiotopic processing. However, as in Experiment 1, there was no transfer of the learned spatial suppression to the spatiotopic location after eye movements. If anything, the data suggests that the learned suppression still persisted in retinotopic coordinates, characterized by a positive error rate slope across the retinotopic towards the spatiotopic location (in line with the scenario in Figure 2A). But in contrast to the findings of Experiment 1, this pattern occurred only for the error rates and not for the RTs. A possible explanation for this discrepancy is that the stimuli were only presented for 150 ms and not until response, making the task very challenging. As a result, participants may have been more inclined to make fast guesses, resulting in less informative reaction times. Experiment 2b addressed this issue by extending the stimulus display duration to 2000 ms or until a response was made. The aim of Experiment 2b was to determine whether the short display time in Experiment 2a was responsible for the effect being observed only in error rates and not in RTs.

## Experiment 2b

### Methods

Experiment 2b was identical to Experiment 2a, except that the stimuli were presented for 2000 ms or until response (as in Experiment 1). The final models to analyze the RTs and error rates in the test array included only by-participant random intercepts in the random-effects structure.

#### Participants

In Experiment 2b, we anticipated the effect size to fall between that of Experiment 1 and Experiment 2a. This expectation was based on the controlled environment of Experiment 1 leading to a higher effect size, and the challenging nature of the task in Experiment 2a resulting in a lower effect size. Thirty-two adults (16 women, mean age: 27.25 years old) were recruited for monetary compensation via the online platform Prolific (www.prolific.co; £9.25).

### Results

One participant was identified as an outlier and replaced based on average RT (M = 1147 ms; See Statistical analysis section of the Methods of Experiment 1 for the exclusion criteria). Exclusion of incorrect trials (9.1%) and data trimming (2.3%) resulted in an overall loss of 11.3% of the trials for the RT analyses and 2.3% of trials for the error rate analyses.

#### Learning array

For the learning array, repeated-measures ANOVAs with within-subjects factor Distractor condition (no distractor, HP location and LP location) yielded a main effect for RTs (*F* (2, 62) = 135.29, *p* < .001, 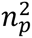 = 0.81) as well as for error rates (*F* (2, 62) = 66.38, *p* < .001, 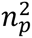 =0.68). Subsequent planned comparisons confirmed that relative to the no distractor condition RTs were longer and error rates were higher at the HP and LP locations (all *t*’s > 6.3, all *p*’s < .001; see Figure 7A and 7C). Crucially, participants were faster (*t* (31) = 6.91, *p* < .001, *d* = 1.22; see Figure 7B) and had lower error rates (*t* (31) = 5.73, *p* < .001, *d* = 1.01; see Figure 7D) when the distractor appeared at the HP location compared to the LP location. Moreover, the RTs and error rates were lowest at the HP location and increased with distance from the HP location (RTs: linear β = 36.35, SE = 5.20, t (12089.22) = 6.99, p < .001; Error rates: linear β = 0.41, SE = 0.066, z = 6.19, p < .001; see Supplementary Materials for figures).

**Figure 7.**
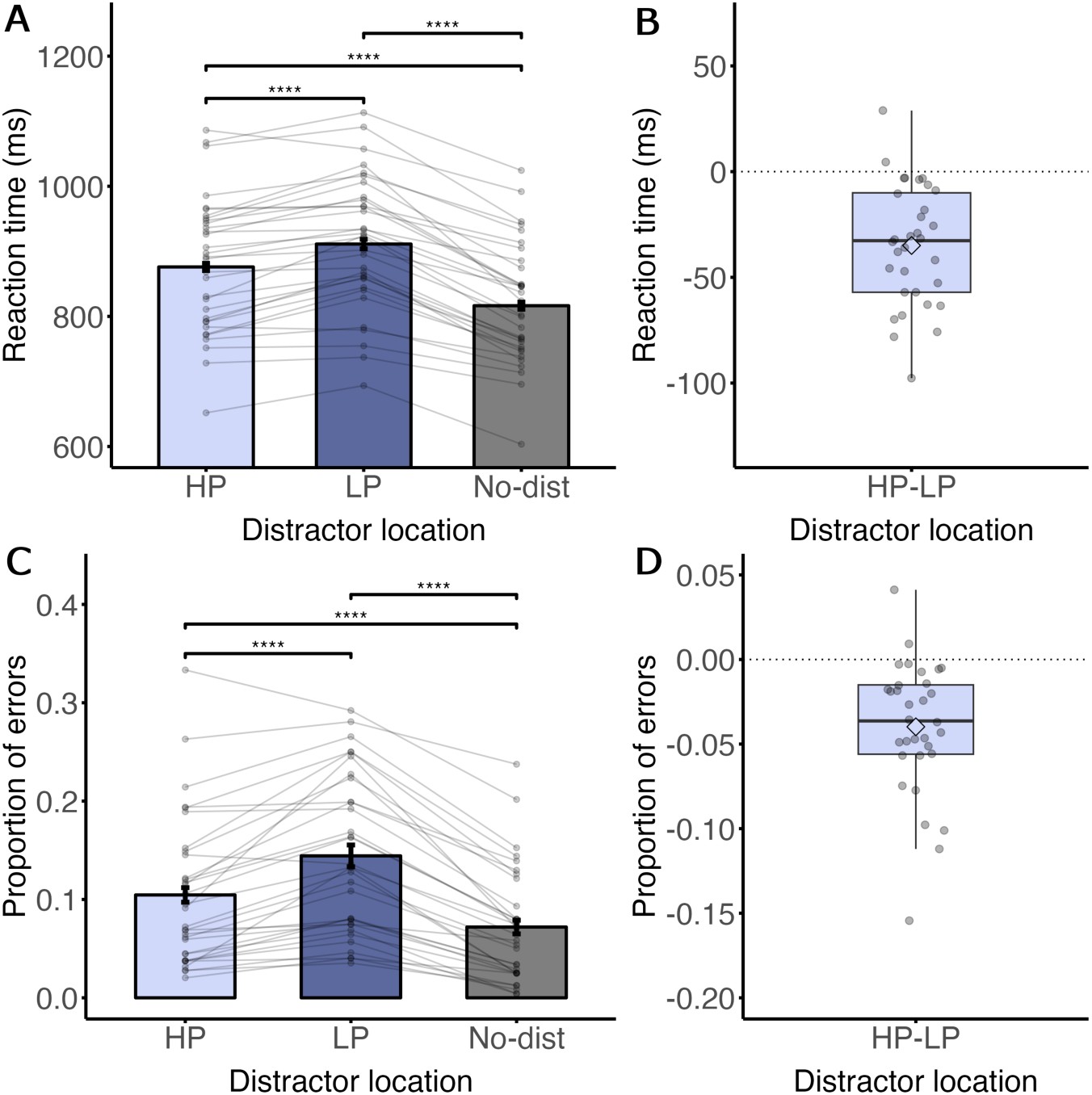
RTs (A and B) and error rates (C and D) in Experiment 2b as a function of distractor location for learning arrays. *Note.* **(A)** RTs in the learning array. The bars show a clear attentional capture effect with longer RTs when the distractor is present. **(B)** The boxplot displays the RT differences between the HP and LP condition in the learning array. Most subjects have shorter RTs when the distractor is presented at the HP location compared to the LP location. **(C)** Error rates in the learning array. The bars demonstrate a pronounced attentional capture effect, indicated by higher error rates when the distractor is present. **(D)** The boxplot displays the error rate differences between the HP and LP condition in the learning array. The majority of subjects exhibit lower error rates when the distractor is presented at the HP compared to the LP location.

#### Test array

Again, we exclusively considered participants who demonstrated a visual statistical learning effect in the learning array for the analyses conducted on the test array (*N* = 30). Counter to the findings of Experiment 2a, as visualized in Figure 8A and 8C respectively, both RT (linear β = 16.53, SE = 7.60, *t* (4402.41) = 2.18, *p* = .030) and error rate (linear β = 0.54, SE = 0.10, *z* = 5.31, *p* < .001) were characterized by a systematic increase across the retinotopic, LP and spatiotopic locations (see Supplementary Materials for parallel analyses involving all participants). Together with the previous experiments these findings show that at least under the present conditions there is no evidence whatsoever that learned spatial suppression is remapped into spatiotopic coordinates following a saccade.

**Figure 8.**
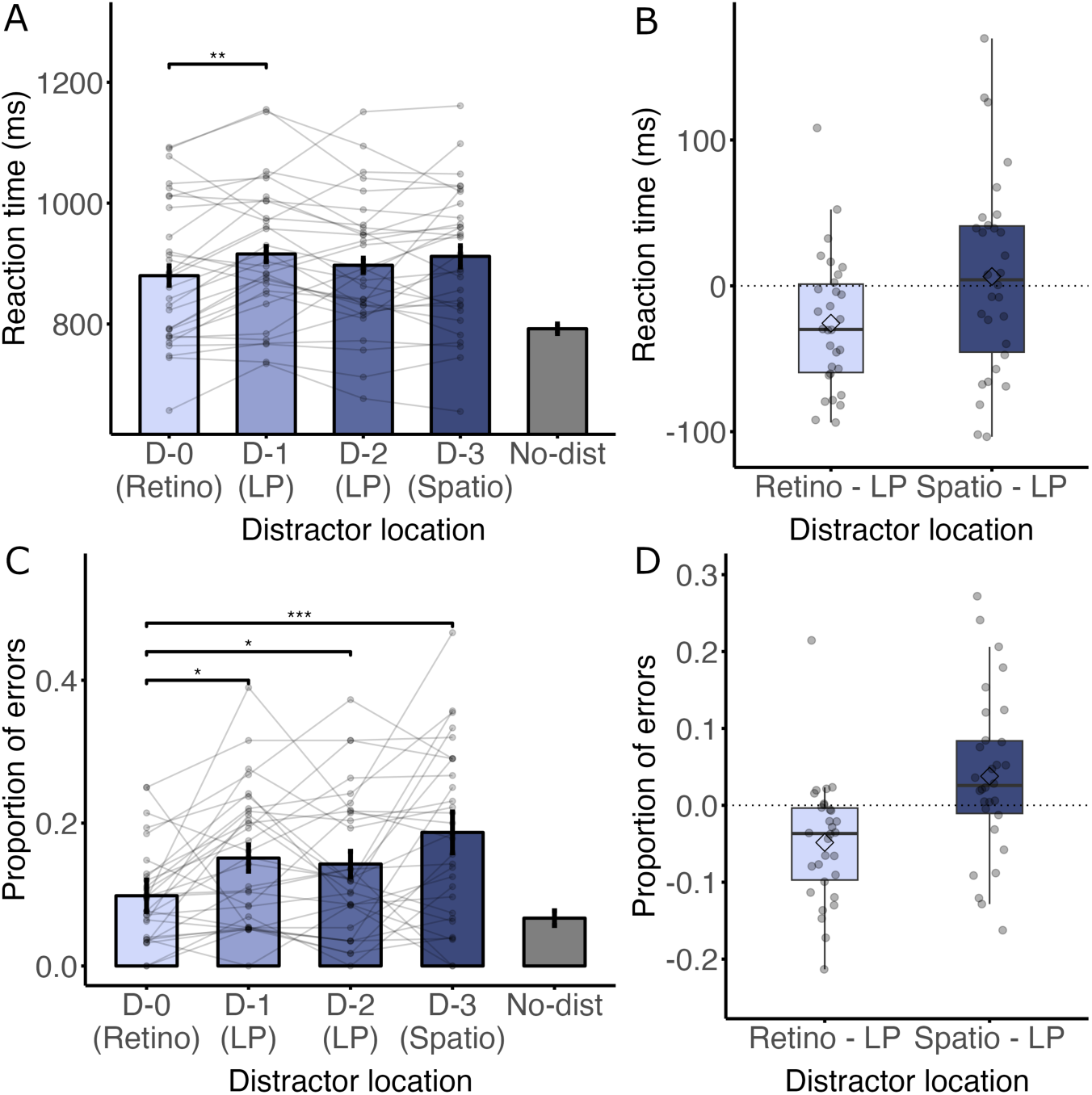
RTs (A and B) and error rates (C and D) in Experiment 2b as a function of distractor location for test arrays. *Note.* Significance bars between the no distractor condition and each distractor location are omitted in the figure (all *t*’s > 2.8 and *p*’s < .05). **(A)** RTs in the test array. Similar to experiment 1, the bars show a positive slope across the retinotopic, LP and spatiotopic locations (linear β = 16.53, SE = 7.60, *t* (4402.41) = 2.18, *p* = .030) **(B)** The boxplot displays the RT differences between the retinotopic and LP location and the spatiotopic and LP location. **(C)** Error rates in the test array. The bars show a systematic increase in error rates across the retinotopic, LP and spatiotopic locations (linear β = 0.54, SE = 0.10, *z* = 5.31, *p* < .001) **(D)** The boxplot displays the error rate differences between the retinotopic and LP location and the spatiotopic and LP location.

### Discussion

Experiment 2b replicated the results of Experiment 1 and demonstrated that, following eye movements, suppression effects due to statistical learning remain in retinotopic coordinates, while there was no transfer of the suppression to spatiotopic coordinates, even when visual landmarks are present to impose a spatial reference frame.

## General Discussion

The present study shows that participants learn the statistical regularities presented in the display and adapt their selection priorities accordingly. More importantly, the current study provides compelling new evidence that the attentional suppression effect due to statistical learning operates in retinotopic coordinates rather than spatiotopic coordinates. Following a saccade to a new location, we see that the same location relative to the eyes is suppressed.

These findings provide some important insight about the underlying mechanism. Given that suppression is only found in retinotopic coordinates, it is possible that learned suppression is resolved by changing synaptic weights in early visual areas, as the initial input to visual cortex is retinotopic. Importantly, it has been suggested that the brain exclusively encodes spatial information within retinotopic maps and does not contain explicit spatiotopic representations (Golomb et al., 2008; Golomb & Kanwisher, 2012; Mathôt & Theeuwes, 2011). Indeed, it has been shown that retinotopy is preserved throughout higher visual areas (Golomb & Kanwisher, 2012). A plausible mechanism for representing topographic maps involves the remapping of retinotopic maps, potentially triggered by eye movement signals, such as a corollary discharge (Wurtz et al., 2011b). Notably, behavioral studies on endogenous attention (Golomb et al., 2008, 2010) and exogenous attention (Mathôt & Theeuwes, 2010b) reveal a gradual remapping of attention from retinotopic to spatiotopic coordinates following eye movements. It has been suggested that the frontal eye field (FEF) is a central source of remapping, with early visual cortices playing a comparatively minor role (Mathôt & Theeuwes, 2011). Given the findings of the current study, the question remains as to why this remapping phenomenon does not seem to apply to the observed suppression effects. This leads to the hypothesis that the suppression effect observed in the current study may be resolved primarily in early visual cortices, without extending to the FEF, in contrast to top-down or bottom-up attentional processes.

Recently, however, a study with a similar design found that spatiotopic suppression can occur under specific conditions (Chang & Golomb, 2024). As in our experiments, in the Chang & Golomb (2024) study there was a learning and test array with overlapping stimulus locations at the center of the screen. Comparable to the current findings, they found that when participants learn the spatial probabilities in the learning array and make a one-time switch in fixation to the test array, the learned suppression remained in retinotopic coordinates. However, when participants dynamically shifted their fixation between the arrays, and the spatial imbalance was maintained, suppression occurred in spatiotopic rather than retinotopic coordinates. They concluded that suppression is generally retinotopic but can be spatiotopic in dynamic contexts.

In our study, participants also frequently varied their fixation, so spatiotopic suppression might have been expected in this context as well. However, the key difference between our study and that of Chang & Golomb (2024) is that their participants varied their fixation from the beginning of the experiment, while our participants started with two blocks of fixation on just one side of the display. A possible explanation for the discrepancy in the results is that, at the start of our experiment, suppressing the retinotopic location was sufficient. However, when the display started dynamically switching between sides, participants were unable to transition to spatiotopic suppression.

Learned suppression is often characterized not only by faster responses and fewer errors when the distractor is at the HP distractor location compared to all other locations, but also by slower responses and more errors when the target is at the HP location (Ferrante et al., 2018; Wang & Theeuwes, 2018c, 2018a, 2018b). In the current study, however, we did not observe this pattern consistently across all experiments (see Supplementary Materials for more detailed results). In Experiment 1, we found no significant differences in RTs or error rates in either the learning or test arrays, though error rates showed marginal trends when the target was at the HP distractor location in the learning array (*p* = .12) and at the retinotopic location in the test array (*p* = .075). In Experiment 2a, increased RTs emerged only at the retinotopic location in the test array (*p* < .001) but not at the HP location in the learning array (*p’s* > .23). In contrast, Experiment 2b showed significantly increased RTs at the HP and retinotopic locations in the learning (*p* < .01) and test arrays (*p* < .001), respectively. It should be noted that the effect on target processing are typically weaker compared to distractor tuned analysis, and sometimes even absent (e.g. van Moorselaar et al., 2021). Given that the current study included only a few trials for the target-related analyses, the low statistical power makes it unsurprising that these effects were not consistently reliable across experiments.

It is important to emphasize that Experiment 2B, in which eye movements were not monitored, served solely as a control experiment. On the basis of this experiment alone, no firm conclusions can be drawn regarding whether the designated retinotopic location consistently aligned with the same retinal position across trials. However, the central findings are conveyed by Experiment 2A, in which brief stimulus durations (<150 ms) were employed specifically to preclude directed eye movements, thereby ensuring the validity of the retinotopic manipulation. In this context, we observed clear retinotopic suppression as reflected in the error rates. Experiment 2B was included only to examine whether a comparable effect would emerge in RT measures.

The fact that participants suppressed the same location relative to the fixation cross, even when they were free to move their eyes during the search, could still be explained by retinotopic suppression, as participants may have kept their fixation across trials. However, another possibility is that suppression relied on a head-centered, egocentric (self-referenced) representation, in line with the findings of Jiang & Swallow (2013, 2014), who demonstrated that attentional biases can shift with the observer’s viewpoint rather than remaining anchored to a specific spatiotopic or retinotopic location.

Specifically, Jiang & Swallow (2013, 2014) conducted a series of experiments demonstrating that attentional enhancement due to probability cuing is dependent on the participants’ viewpoint. Participants were tasked with locating a T among L’s displayed on a tablet mounted on a stand. Unbeknownst to the participants, the target appeared more frequently in one quadrant compared to the others. As expected, the study revealed an attentional bias towards the quadrant that was likely to contain the target. However, intriguingly, when participants moved around the tablet, the attentional facilitation appeared to move along with the participant’s viewpoint rather than remaining in the spatiotopic location. It is important to note that in these experiments, each trial began with a fixation dot randomly placed within a central region and participants were allowed to freely move their eyes during search.

This implies that the likely target location was not learned in a retinotopic manner (i.e., relative to the eyes) but within an egocentric reference frame (i.e., relative to the head-body). Consequently, it appears that there is not only a lack of remapping of statistical learning effects from retinotopic to spatiotopic coordinates following eye movements but also an absence of updating the egocentric reference frame to an environmentally stable reference frame after body and head movements (but also see Jiang et al., 2014; Smith et al., 2010; Zheng et al., 2021). Given our continuous eye and body movements, the practical use of learned attentional biases becomes uncertain when they are not remapped from retinotopic or egocentric coordinates to spatiotopic coordinates.

An alternative explanation for our findings, which we have not yet explored, is that suppression might not rely on either retinotopic or spatiotopic coordinates but instead on an object-based coordinate system. Although we attempted to create a stable environmental context using placeholders and a grid, the six colored stimuli surrounding the fixation point could have functioned as a single object. Therefore, the suppression effect may have been based on object-based coordinates rather than retinotopic coordinates. Indeed, it has been shown that attentional enhancement driven by statistical learning can also occur within objects, irrespective of the object’s location or orientation in space (Ilksoy et al., 2025; van Moorselaar & Theeuwes, 2023, 2024). Additionally, it has been shown that habituation to abrupt distractor onsets is susceptible to target-distractor configurations (Turatto et al., 2024).

As a result, in the current design we cannot definitively distinguish between retinotopic and object-based suppression. However, we can only conclude that, in this context, suppression was not remapped to spatiotopic coordinates.

In summary, the findings of the current study indicate that, following saccadic eye movements, suppression effects persist in retinotopic coordinates, with no observed transfer of suppression to spatiotopic coordinates. Further research is needed to determine whether situations exist in which implicit attentional biases are remapped to environmentally stable coordinates.

