## Supplementary Materials for "Distractor suppression operates exclusively in retinotopic coordinates"

**Supplementary table 1**

Mean performance of all participants (and not only the participants that exhibited visual statistical learning) in the test array across Experiments 1, 2a, and 2b

| Condition | Reaction time (ms) | | | Proportion of errors | | |
| --- | --- | --- | --- | --- | --- | --- |
|  | n | M | CI^a^ | n | M | CI^a^ |
| Experiment 1 |  |  |  |  |  |  |
| D-0 (Retino) | 706 | 752 | 13.8 | 755 | 0.065 | 0.020 |
| D-1 (LP) | 1424 | 768 | 10.3 | 1565 | 0.091 | 0.016 |
| D-2 (LP) | 1431 | 772 | 10.6 | 1563 | 0.084 | 0.015 |
| D-3 (Spatio) | 705 | 781 | 14.4 | 772 | 0.087 | 0.022 |
| Experiment 2a |  |  |  |  |  |  |
| D-0 (Retino) | 1049 | 741 | 9.0 | 1442 | 0.27 | 0.025 |
| D-1 (LP) | 2057 | 726 | 6.2 | 2899 | 0.29 | 0.018 |
| D-2 (LP) | 2039 | 723 | 6.4 | 2909 | 0.30 | 0.018 |
| ` | 940 | 740 | 9.7 | 1435 | 0.34 | 0.027 |
| Experiment 2b |  |  |  |  |  |  |
| D-0 (Retino) | 827 | 888 | 15.8 | 926 | 0.11 | 0.022 |
| D-1 (LP) | 1557 | 914 | 12.0 | 1840 | 0.15 | 0.018 |
| D-2 (LP) | 1584 | 899 | 11.6 | 1848 | 0.14 | 0.017 |
| D-3 (Spatio) | 748 | 914 | 17.0 | 920 | 0.19 | 0.028 |

^a^ 95% within-subject confidence interval

**Supplementary table 2**

(G)LMM results of RT and error rate analyses for all participants (and not only the participants that exhibited visual statistical learning) in the test array across Experiments 1, 2a, and 2b

| Measure | Fixed effect^a^ | Estmate (β) | SE | df | t/z | p-value | p < .05 |
| --- | --- | --- | --- | --- | --- | --- | --- |
| Experiment 1 |  |  |  |  |  |  |  |
| Reaction time | Linear | 19.78 | 8.32 | 22.79 | 2.38 | .026 | * |
| Error rate | Linear | 0.19 | 0.14 |  | 1.41 | 0.16 | n.s. |
| Error rate | Quadratic | -0.24 | 0.19 |  | 1.28 | 0.20 | n.s. |
| Experiment 2a |  |  |  |  |  |  |  |
| Reaction time | Quadratic | 27.57 | 6.94 | 194.51 | 3.97 | < .001 | *** |
| Error Rate | Linear | 0.27 | 0.085 |  | 3.20 | < .01 | ** |
| Experiment 2b |  |  |  |  |  |  |  |
| Reaction time | Linear | 12.97 | 7.37 | 4684.47 | 1.76 | 0.079 | . |
| Error rate | Linear | 0.49 | 0.097 |  | 5.067 | < .001 | *** |

*Note.* Significant effects are marked as follows: *p < .05, **p < .01, ***p < .001, and non-significant results are indicated by "n.s."

*^a^* The data were fitted to both linear and quadratic models. Only the statistically significant model is shown in the table. If neither model is significant, results for both insignificant models are presented.

**Supplementary table 3**

Repeated-measures ANOVA results for the test array across Experiments 1, 2a, and 2b

| Measure | df | F | p-value | p < .05 | Partial η^2^ |
| --- | --- | --- | --- | --- | --- |
| Experiment 1 |  |  |  |  |  |
| Reaction time | 4, 84 | 22.32 | < .001 | *** | 0.52 |
| Error rate | 4, 84 | 6.44 | < .001 | *** | 0.24 |
| Experiment 2a |  |  |  |  |  |
| Reaction time | 4, 148 | 18.32 | < .001 | *** | 0.33 |
| Error rate | 3.24, 120.05 | 26.26 | < .001 | *** | 0.42 |
| Experiment 2b |  |  |  |  |  |
| Reaction time | 3.04, 88.08 | 32.45 | < .001 | *** | 0.53 |
| Error rate | 2.76, 80.05 | 15.62 | < .001 | *** | 0.35 |

*Note.* Significant effects are marked as follows: *p < .05, **p < .01, ***p < .001, and non-significant results are indicated by "n.s.".

**Supplementary table 4**

Planned paired-sample t-test results for the test array across Experiments 1, 2a, and 2b

| Measure | Planned comparison | df | t-value | p-value | p_holm_ | p_holm_ < .05 | Cohen’s d |
| --- | --- | --- | --- | --- | --- | --- | --- |
| Experiment 1 |  |  |  |  |  |  |  |
| Reaction time |  |  |  |  |  |  |  |
|  | D-0 (Retino) vs. D-1 (LP) | 21 | 2.15 | .044 | 0.13 | n.s. | 0.46 |
|  | D-0 (Retino) vs. D-2 (LP) | 21 | 2.39 | .026 | 0.10 | n.s. | 0.51 |
|  | D-0 (Retino) vs. D-3 (Spatio) | 21 | 3.026 | < .01 | 0.030 | * | 0.65 |
|  | D-3 (Spatio) vs. D-1 (LP) | 21 | 1.24 | .23 | 0.46 | n.s. | 0.26 |
|  | D-3 (Spatio) vs. D-2 (LP) | 21 | 0.85 | .41 | 0.46 | n.s. | 0.18 |
| Error rate |  |  |  |  |  |  |  |
|  | D-0 (Retino) vs. D-1 (LP) | 21 | 3.78 | < .01 | < .01 | ** | 0.81 |
|  | D-0 (Retino) vs. D-2 (LP) | 21 | 3.14 | < .01 | 0.030 | * | 0.67 |
|  | D-0 (Retino) vs. D-3 (Spatio) | 21 | 2.16 | .043 | 0.17 | n.s. | 0.46 |
|  | D-3 (Spatio) vs. D-1 (LP) | 21 | 0.58 | .57 | 1 | n.s. | 0.12 |
|  | D-3 (Spatio) vs. D-2 (LP) | 21 | 0.15 | .89 | 1 | n.s. | 0.031 |
| Experiment 2a |  |  |  |  |  |  |  |
| Reaction time |  |  |  |  |  |  |  |
|  | D-0 (Retino) vs. D-1 (LP) | 37 | 1.87 | 0.07 | 0.21 | n.s. | 0.30 |
|  | D-0 (Retino) vs. D-2 (LP) | 37 | 1.49 | 0.15 | 0.29 | n.s. | 0.24 |
|  | D-0 (Retino) vs. D-3 (Spatio) | 37 | 1.29 | 0.21 | 0.29 | n.s. | 0.21 |
|  | D-3 (Spatio) vs. D-1 (LP) | 37 | 3.25 | < .01 | 0.01 | * | 0.53 |
|  | D-3 (Spatio) vs. D-2 (LP) | 37 | 3.35 | < .01 | 0.01 | * | 0.54 |
| Error rate |  |  |  |  |  |  |  |
|  | D-0 (Retino) vs. D-1 (LP) | 37 | 1.03 | 0.31 | 0.53 | n.s. | 0.17 |
|  | D-0 (Retino) vs. D-2 (LP) | 37 | 1.13 | 0.27 | 0.53 | n.s. | 0.18 |
|  | D-0 (Retino) vs. D-3 (Spatio) | 37 | 2.42 | 0.020 | 0.10 | n.s. | 0.39 |
|  | D-3 (Spatio) vs. D-1 (LP) | 37 | 2.05 | 0.047 | 0.19 | n.s. | 0.33 |
|  | D-3 (Spatio) vs. D-2 (LP) | 37 | 1.74 | 0.090 | 0.27 | n.s. | 0.28 |
| Experiment 2b |  |  |  |  |  |  |  |
| Reaction time |  |  |  |  |  |  |  |
|  | D-0 (Retino) vs. D-1 (LP) | 29 | 3.54 | < .01 | < .01 | ** | 0.65 |
|  | D-0 (Retino) vs. D-2 (LP) | 29 | 1.70 | 0.10 | 0.30 | n.s. | 0.31 |
|  | D-0 (Retino) vs. D-3 (Spatio) | 29 | 2.21 | 0.035 | 0.14 | n.s. | 0.40 |
|  | D-3 (Spatio) vs. D-1 (LP) | 29 | 0.18 | 0.86 | 0.86 | n.s. | 0.033 |
|  | D-3 (Spatio) vs. D-2 (LP) | 29 | 1.40 | 0.17 | 0.35 | n.s. | 0.25 |
| Error rate |  |  |  |  |  |  |  |
|  | D-0 (Retino) vs. D-1 (LP) | 29 | 3.18 | < .01 | 0.020 | * | 0.58 |
|  | D-0 (Retino) vs. D-2 (LP) | 29 | 2.83 | < .01 | 0.032 | * | 0.52 |
|  | D-0 (Retino) vs. D-3 (Spatio) | 29 | 4.55 | < .001 | < .001 | *** | 0.83 |
|  | D-3 (Spatio) vs. D-1 (LP) | 29 | 1.49 | 0.15 | 0.15 | n.s. | 0.27 |
|  | D-3 (Spatio) vs. D-2 (LP) | 29 | 2.31 | 0.028 | 0.056 | . | 0.42 |

*Note.* Significant effects are marked as follows: *p < .05, **p < .01, ***p < .001, and non-significant results are indicated by "n.s.".

**Supplementary table 5**

Performance as a function of target location on distractor absent trials across Experiment 1, 2a and 2b

| Condition | Reaction time (ms) | | | Proportion of errors | | |
| --- | --- | --- | --- | --- | --- | --- |
|  | n^a^ | M | CI^b^ | n^a^ | M | CI^b^ |
| Experiment 1 |  |  |  |  |  |  |
| Learning array |  |  |  |  |  |  |
| HP dist. loc | 1336 | 734 | 14.5 | 1424 | 0.062 | 0.018 |
| LP dist. loc | 6514 | 737 | 6.2 | 6850 | 0.049 | 0.0072 |
| Test array |  |  |  |  |  |  |
| D-0 (Retino) | 333 | 701 | 20.8 | 357 | 0.066 | 0.030 |
| D-1 (LP) | 686 | 702 | 12.7 | 721 | 0.048 | 0.018 |
| D-2 (LP) | 699 | 695 | 12.9 | 722 | 0.033 | 0.015 |
| D-3 (Spatio) | 354 | 701 | 19.7 | 370 | 0.043 | 0.024 |
| Experiment 2a |  |  |  |  |  |  |
| Learning array |  |  |  |  |  |  |
| HP | 1563 | 708 | 8.6 | 1886 | 0.17 | 0.023 |
| LP | 8353 | 704 | 3.7 | 9942 | 0.16 | 0.010 |
| Test array |  |  |  |  |  |  |
| D-0 (Retino) | 455 | 701 | 12.6 | 531 | 0.15 | 0.034 |
| D-1 (LP) | 1021 | 700 | 8.1 | 1193 | 0.14 | 0.023 |
| D-2 (LP) | 1053 | 692 | 7.6 | 1192 | 0.12 | 0.021 |
| D-3 (Spatio) | 428 | 675 | 11.8 | 508 | 0.15 | 0.036 |
| Experiment 2b |  |  |  |  |  |  |
| Learning array |  |  |  |  |  |  |
| HP | 1108 | 840 | 16.7 | 1212 | 0.086 | 0.022 |
| LP | 5958 | 813 | 6.8 | 6400 | 0.069 | 0.0086 |
| Test array |  |  |  |  |  |  |
| D-0 (Retino) | 377 | 829 | 23.7 | 409 | 0.078 | 0.029 |
| D-1 (LP) | 891 | 788 | 13.0 | 948 | 0.060 | 0.017 |
| D-2 (LP) | 880 | 785 | 13.0 | 942 | 0.066 | 0.018 |
| D-3 (Spatio) | 385 | 781 | 21.6 | 415 | 0.072 | 0.028 |

*Note.* Table reflects data from those participants who exhibited a visual statistical learning effect in the learning array (same participants as in the main analyses).

^a^ The means for the D-0 and D-3 conditions are based on approximately 15 trials per participant resulting in reduced statistical power in comparison to the main analyses.

^b^ 95% within-subject confidence interval.

**Supplementary table 6**

Planned paired t-tests on the RTs and error rates as a function of target location on distractor absent trials in the learning array across Experiments 1, 2a, and 2b

| Measure | Planned comparison | df | t-value | p-value | p < .05 | Cohen’s d |
| --- | --- | --- | --- | --- | --- | --- |
| Experiment 1 |  |  |  |  |  |  |
| Reaction time | HP dist. loc vs. LP dist. loc | 23 | 0.17 | .87 | n.s. | 0.035 |
| Error rate | HP dist. loc vs. LP dist. loc | 23 | 1.62 | .12 | n.s. | 0.33 |
| Experiment 2a |  |  |  |  |  |  |
| Reaction time | HP dist. loc vs. LP dist. loc | 49 | 1.22 | .23 | n.s. | 0.17 |
| Error rate | HP dist. loc vs. LP dist. loc | 49 | 1.15 | .26 | n.s. | 0.16 |
| Experiment 2b |  |  |  |  |  |  |
| Reaction time | HP dist. loc vs. LP dist. loc | 31 | 3.44 | < .01 | ** | 0.61 |
| Error rate | HP dist. loc vs. LP dist. loc | 31 | 1.50 | .15 | n.s. | 0.26 |

*Note.* Significant effects are marked as follows: *p < .05, **p < .01, ***p < .001, and non-significant results are indicated by "n.s.".

**Supplementary table 7**

(G)LMM results of RTs and error rates as a function of target location on distractor absent trials in the test array across of Experiments 1, 2a, and 2b

| Measure | Fixed effect^a^ | Estmate (β) | SE | df | t/z | p-value | p < .05 |
| --- | --- | --- | --- | --- | --- | --- | --- |
| Experiment 1 |  |  |  |  |  |  |  |
| Reaction time | Linear | -1.26 | 8.16 | 2051.16 | 0.15 | .88 | n.s. |
| Reaction time | Quadratic | 12.23 | 11.73 | 2051.025 | 1.042 | .30 | n.s. |
| Error rate | Linear | -0.42 | 0.24 |  | 1.78 | .075 | . |
| Error rate | Quadratic | 0.67 | 0.40 |  | 1.70 | .090 | . |
| Experiment 2a |  |  |  |  |  |  |  |
| Reaction time | Linear | -18.37 | 5.28 | 2919.49 | 3.48 | < .001 | *** |
| Error rate | Linear | -0.026 | 0.12 |  | 0.21 | .83 | n.s. |
| Error Rate | Quadratic | 0.32 | 0.19 |  | 1.67 | .096 | . |
| Experiment 2b |  |  |  |  |  |  |  |
| Reaction time | Linear | -31.52 | 8.89 | 2503.15 | 3.55 | < .001 | *** |
| Error rate | Linear | -0.023 | 0.19 |  | 0.12 | .90 | n.s. |
| Error rate | Quadratic | 0.31 | 0.30 |  | 1.035 | .30 | n.s. |

*Note.* Significant effects are marked as follows: *p < .05, **p < .01, ***p < .001, and non-significant results are indicated by "n.s.".

*^a^* The data were fitted to both linear and quadratic models. Only the statistically significant model is shown in the table. If neither model is significant, results for both insignificant models are presented.

**Supplementary figure 1**

*RTs (A) and error rates (B) in Experiment 1 as a function of distractor location for learning arrays.*

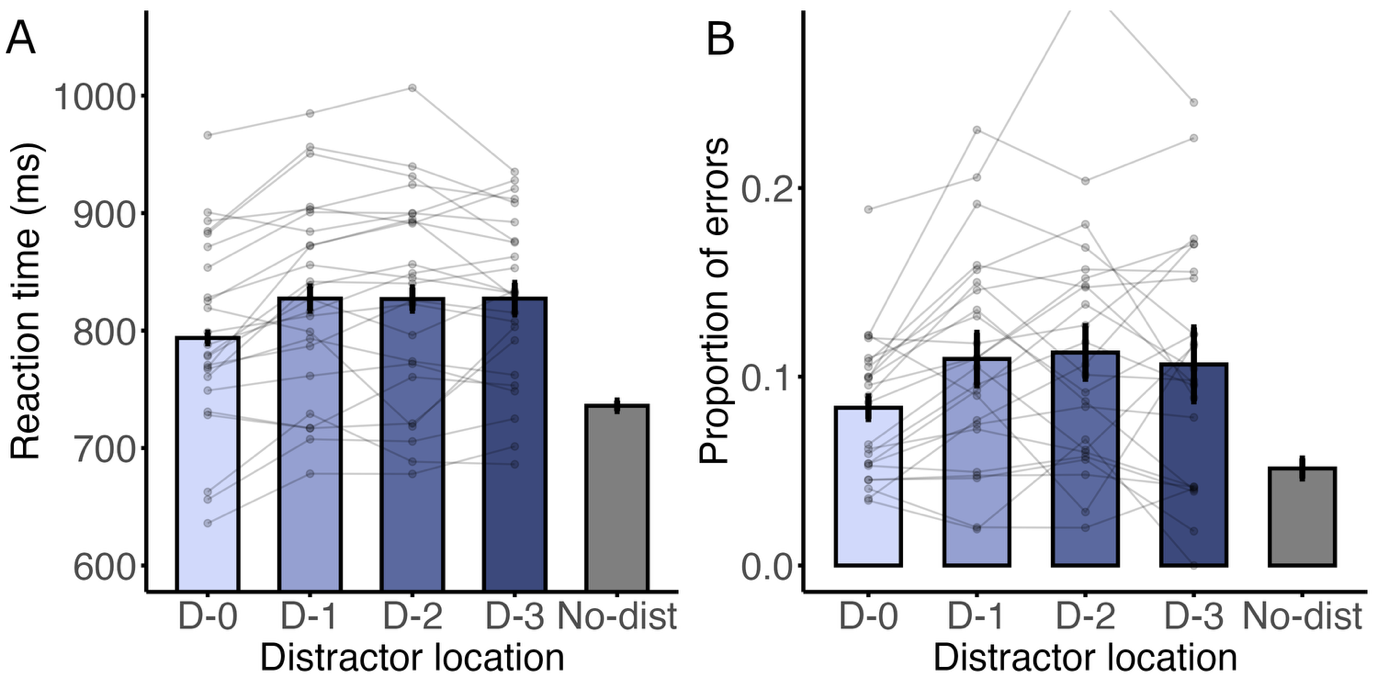

Note. The “D” labels represent the distance to the HP location, with D-0 represeting the HP location itself.

**Supplementary figure 2**

*RTs (A) and error rates (B) in Experiment 2a as a function of distractor location for learning arrays.*

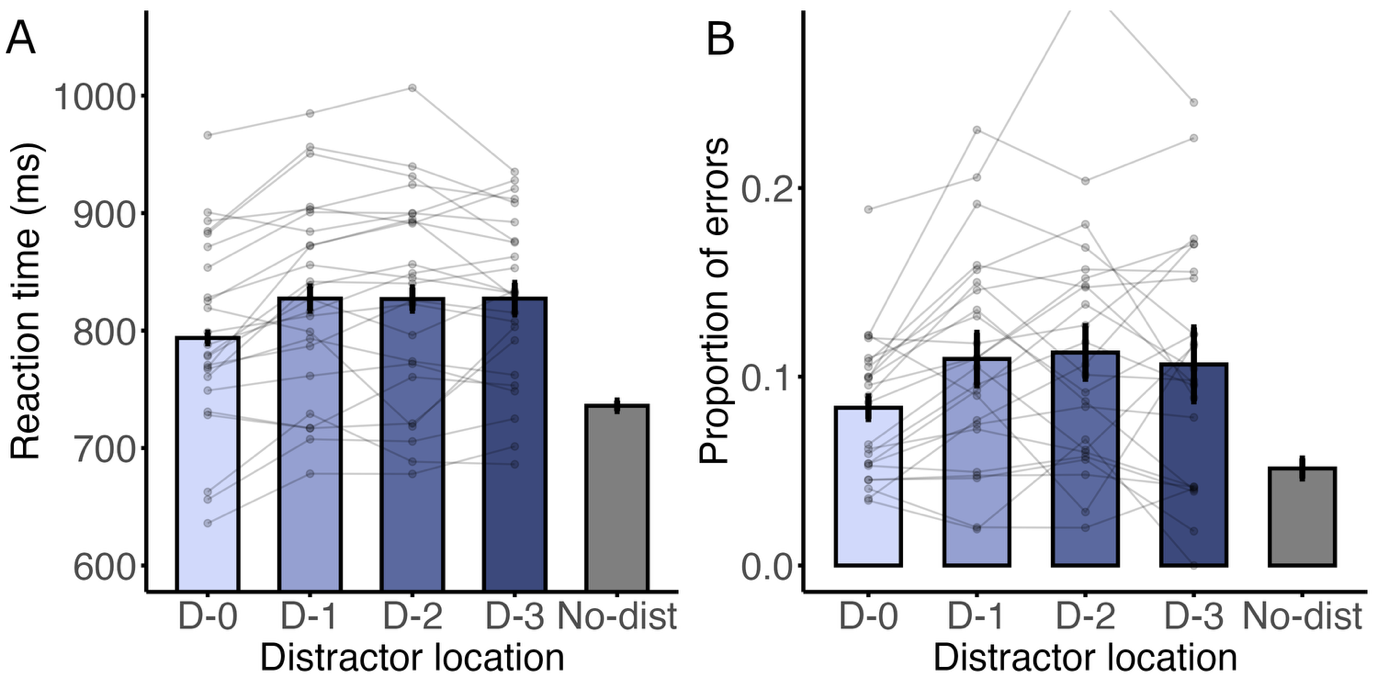

Note. The “D” labels represent the distance to the HP location, with D-0 represeting the HP location itself.

**Supplementary figure 3**

*RTs (A) and error rates (B) in Experiment 2b as a function of distractor location for learning arrays.*

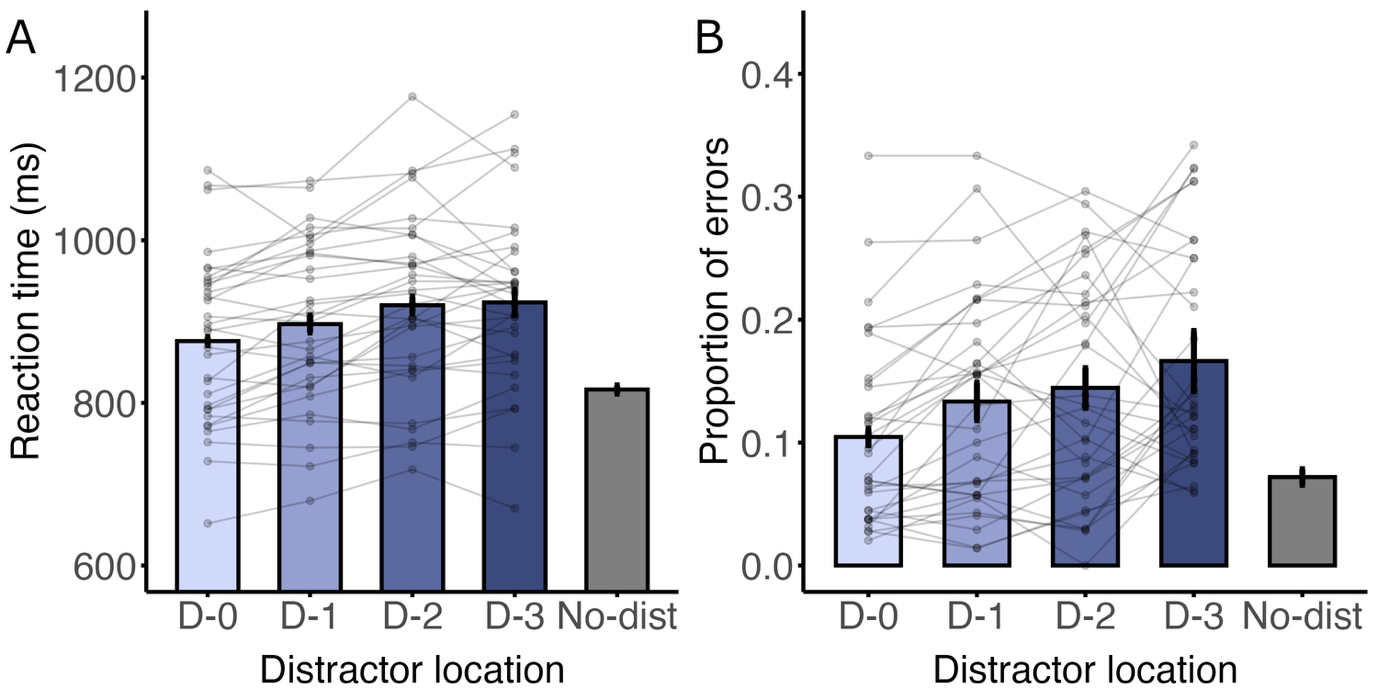

Note. The “D” labels represent the distance to the HP location, with D-0 represeting the HP location itself.
